# Function of *ex vivo* stimulated Natural killer T cells from patients with chronic HIV-1 infection

**DOI:** 10.1101/831834

**Authors:** Prakash Thapa, Pramod Nehete, Hong He, Bharti Nehete, Stephanie J. Buchl, Lara M. Bull, Hongzhou Lu, Roberto C. Arduino, K. Jagannadha Sastry, Dapeng Zhou

**Affiliations:** Department of Melanoma Medical Oncology, The University of Texas M.D. Anderson Cancer Center, Houston, TX; Department of Veterinary Sciences, The University of Texas M.D. Anderson Cancer Center, Houston, TX; Department of Immunology, The University of Texas M.D. Anderson Cancer Center, Houston, TX; Clinical Epidemiology and Outcomes Program, Baylor College of Medicine, Houston, TX; Shanghai Public Health Clinical Center, Shanghai, China; Division of Infectious Diseases, Department of Medicine, The University of Texas Medical School at Houston, Houston, TX; Graduate School of Biological Sciences, The University of Texas Health Science Center at Houston, Houston, TX; School of Medicine, Tongji University, Shanghai, China

**Keywords:** natural killer T cells, alpha-galactosylceramide, dendritic cells, HIV-1, innate immunity

## Abstract

Natural killer T (NKT) cells are innate immune cells that are responsible for the first line of antiviral defense, through crosstalk with downstream antigen-presenting cells, natural killer cells, and adaptive immune cells. Previous studies have indicated that NKT cell function is severely impaired in patients with chronic HIV-1 infection. It was reported that alpha-galactosylceramide, a potent agonist antigen for NKT cells, failed to trigger the expansion of NKT cells, or the production of anti-viral cytokines by NKT cells from HIV-1 infected patients in an in vitro assay, in which peripheral blood mononuclear cells (PBMCs) were cultured in the presence of alpha-galactosylceramide. In this study, we stimulated banked peripheral blood mononuclear cells from HIV-1-infected patients with dendritic cells (DC) generated ex vivo and loaded with alpha-galactosylceramide. The results showed that NKT cells were expanded in HIV infected subjects except in patients with advanced AIDS. Expanded NKT cells were capable of producing antiviral cytokines. Our results indicate that NKT cells in HIV infected individuals are potential targets for therapeutic intervention.

## Introduction

Reconstitution of the immune system in patients with chronic HIV-1 infection is important not only for eliminating the virus and virus-infected cells, but also for preventing the onset of cancers and opportunistic infections due to a virus-induced loss of CD4 T cells. A recently discovered class of innate immune cells, named natural killer T (NKT) cells, may “jump-start” antiretroviral immune responses through their unique ability to activate antigen-presenting cells (e.g., dendritic cells [DCs]), natural killer (NK) cells, and adaptive immune cells including CD8 T cells (1–2). The regulatory effects of NKT cells on DCs, NK cells, and CD8 cells are mediated by direct cell-to-cell contact and the large amount of cytokines that NKT cells produce. NKT cells include both CD4+ and CD4-subsets. The CD4+ NKT cells are susceptible to HIV infection, while susceptibility of CD4-NKT cells remains controversial (3–5). After antiretroviral therapy, CD4-NKT cells could be recovered from HIV infected patients (i.e., increases in CD4-NKT cell populations were noted) (6). Furthermore, during acute HIV-1 infection, impaired NKT cell function could be improved by effective highly active antiretroviral therapy (7). Moll et al. reported that when immuno-stimulatory reagents such as interleukin-2 (IL-2) were combined with antiviral therapy, both CD4+ and CD4-NKT cells were recovered (8). However, it remains unclear whether the recovered NKT cells are functional. Using results from an in vitro culture assay, Moll et al recently suggested that NKT cell function in patients with chronic HIV-1 infection could not be restored after antiretroviral treatment (9).

In contrast to conventional T cells, which recognize peptide antigens presented by MHC molecules, NKT cells recognize glycolipid antigens presented by a non-MHC and non-polymorphic antigen presenting molecule, CD1d. One potent agonist antigen for NKT cells is alpha-galactosylceramide (10), a glycolipid that is currently being tested as an immuno-modifier/enhancer to treat cancer (11, 12) and hepatitis (13, 14). However, the effect of this drug on NKT cells in HIV-infected patients remains unclear. Moll et al reported that even in the presence of IL2 and antiretroviral drugs, alpha-galactosylceramide failed to trigger the expansion of NKT cells, or the production of antiviral cytokines by NKT cells from HIV-1 infected patients in an in vitro assay, in which peripheral blood mononuclear cells (PBMCs) were cultured in the presence of alpha-galactosylceramide (9).

A major limitation to the pharmaceutical use of soluble alpha-galactosylceramide is that alpha-galactosylceramide causes NKT cell anergy after a single dose, because alpha-galactosylceramide can be presented by CD1d-expressing B cells in the peripheral blood and stimulate the NKT cells without proper costimulatory molecules (15–17). To overcome this problem, Dhodapkar and Steinmann developed a cell therapy approach by intravenously injecting alpha-galactosylceramide loaded DCs generated in vitro from patients’ peripheral blood (18). This method prevented NKT cell anergy and showed efficacy for eliciting antigen-specific T cell responses toward cytomegalovirus in cancer patients. Thus therapeutic activation of NKT cells can establish the link between the innate and adaptive arms of immunity and induce recall responses by memory T cells. In this study, we examined whether the function of HIV-infected NKT cells could be reconstituted by stimulation with alpha-galactosylceramide loaded DCs, by in vitro culture experiments using HIV-1-infected NKT cells from patients.

## Methods

### Defined population

Banked frozen PBMCs from antiretroviral therapy naïve, newly diagnosed African American male HIV-1 infected patients were provided by the Clinical Research Core at the Baylor-UT-Houston Center for AIDS Research. We studied PBMCs from the following 3 groups of patients:

1. Low viral load: <50,000 copies/ml, CD4 cell count >350 cells/mm^3^
2. Medium viral load: >50,000 copies/ml, CD4 cell count 50-350 cells/mm^3^
3. High viral load (advanced AIDS): CD4 cell count<50 cells/mm^3^

For each group, PBMC samples from 5 patients were included; in addition, PBMC samples from 5 healthy donors (Gulf Coast Regional Blood Center, Houston, TX) were used as controls, for a total of 20 samples. We used about 10 million of PBMCs purified from 10 ml of peripheral blood, for both HIV infected patients and healthy individuals. All studies were approved by Committee for the Protection of Human Subjects at the University of Texas Health Science Center at Houston with IRB approval and informed patient consent.

### Generation of alpha-galactosylceramide-loaded DCs for in vitro NKT stimulation

alphagalactosylceramide-loaded DCs were produced according to our published protocols (19). Briefly, monocytes were attached to 75-mm Falcon cell culture flasks (BD Sciences) for 2 hours and cultured in Iscove’s modified Dulbecco’s medium (Invitrogen) with 2% human serum (Valley Biomedical Products & Services) with 800 U/mL granulocyte-macrophage colony-stimulating factor (R&D Systems) and 1000 U/mL IL4 (R&D Systems) for 5 days. On day 6 of culture, 100 ng/ml alpha-galactosylceramide (20) was added to the culture medium. At day 6 of culture, 10 ng/ml IL-1β, 10 ng/ml tumor necrosis factor-alpha (TNF-α), 15 ng/ml IL-6 (all from R&D Systems) and 1 μg/ml prostaglandin E2 (EMD Chemicals) were added to trigger the maturation of DCs. The DCs were harvested on day 8 of culture to be used for the stimulation of NKT cells in vitro.

Allogenic DCs generated from healthy donors were used as the antigen presenting cells for alpha-galactosylceramide. We chose to use allogenic DCs for 2 reasons. First, the alpha-galactosylceramide is presented by CD1d molecule, which is non-polymorphic (1–2). Second, it was technically impractical for us to generate autologous DCs from HIV patients, since our study was based on frozen samples in tissue bank, which allows us to have access to only a few million of PBMCs.

### In vitro expansion of NKT cells from PBMCs of HIV patients

We mixed 10 million of PBMCs from HIV patients with 1 million of alpha-galactosylceramide-loaded DCs, and co-cultured them in AIM-V medium (Invitrogen) with 10% human serum (Valley Biomedical Products & Services), 1 μg/ml azidodeoxythymidine (Sigma), and 10 ng/mL IL-15 (R&D Systems). On day 2 of co-culture, 100 U/ml IL2 (Chiron) was added. On day 14 day of co-culture, NKT cells were restimulated by alpha-galactosylceramide-loaded DCs. Expanded NKT cells were quantified by staining with PBS57-CD1d tetramer (NIH Tetramer Facility at Emory University), anti-Vα24 antibody (Beckman Coulter), and anti-CD3 and anti-CD4 antibodies (eBioscience).

### Study on the function of expanded NKT cells

Expanded NKT cells were studied for their capacity to produce antiviral cytokines, by intracellular staining for INF-γ, TNF-α, and IL4 (all from eBioscience) within 12 hours of being stimulated by alphagalactosylceramide-loaded DCs in the presence of Golgi-plug according to the manufacturer’s instructions (BD Biosceinces).

### Statistic analysis

Descriptive statistics (mean, SD, median, minimum and maximum) were calculated for the outcome variables by groups. Distribution of the outcome variables was examined to choose appropriate analysis method. One-way ANOVA tests were applied to test the differences among three groups for each outcome and cell type respectively. All analyses were performed using SAS 9.2.

## Results

### Alpha-galactosylceramide/DC triggered expansion of NKT cells from HIV-1 infected patients

After 2 rounds of stimulating with alpha-galactosylceramide-loaded, monocyte-devied DCs, we observed expansion of NKT cells in the PBMC samples of HIV-1 infected patients with low and medium viral loads (Figure 1). Since we used allogenic DCs to stimulate NKT cell growth, we also observed allogenic expansion of CD4 and CD8 T cells. Thus we could not use the percentage of NKT cells among total T cells as a parameter to measure NKT cell growth. Instead, we used the absolute number of PBS57/CD1d tetramer-positive cells. After the first round of stimulation, NKT cells were expanded by a factor of 10-10^4^ in patients with low and medium viral loads, similar to that of healthy individuals. After second round of stimulaton, however, the patients with medium viral load showed less (subjects MVL4 and MVL5) or no expansion (MVL2 and MVL3) as compared to healthy individuals.

**Figure 1.**
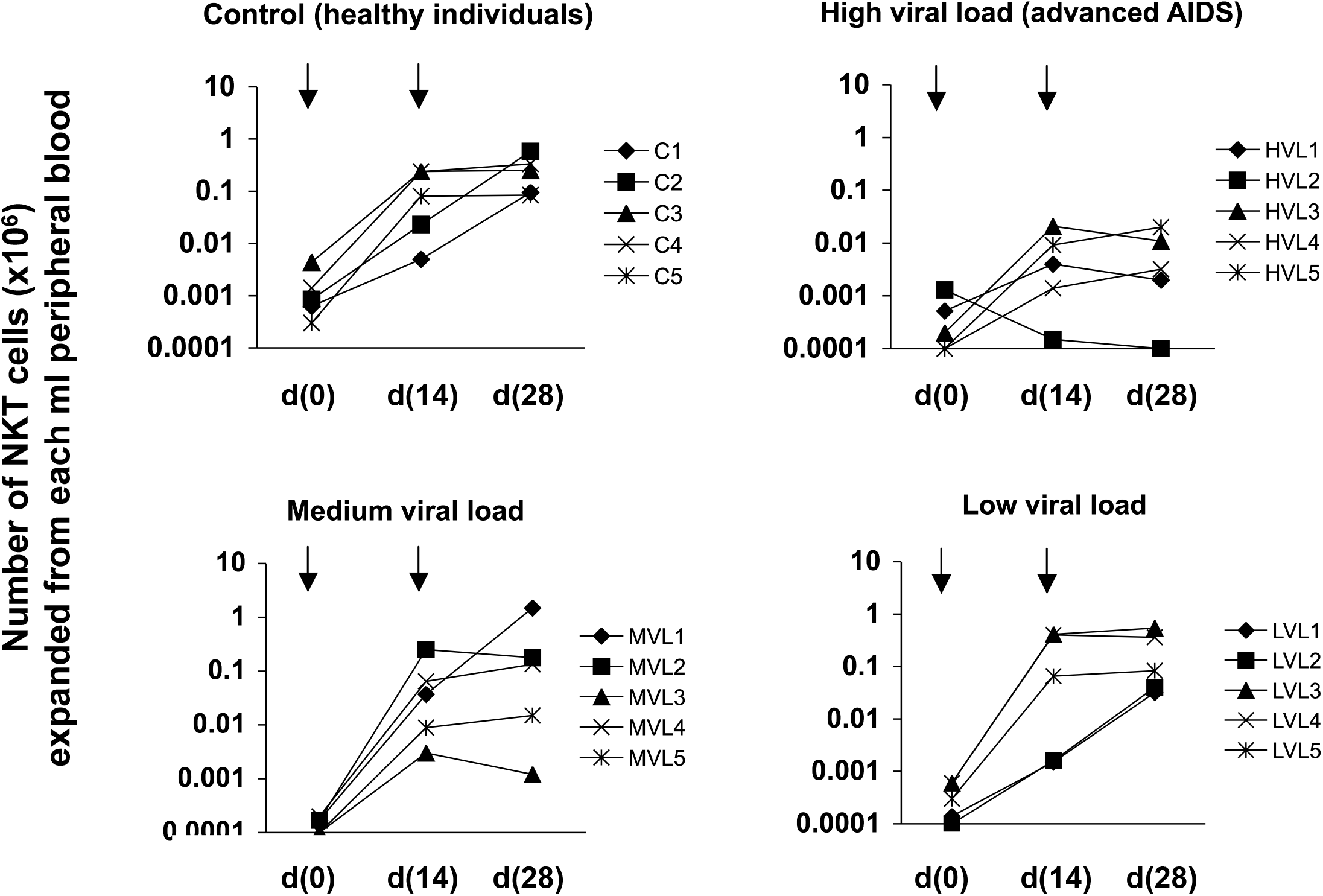
Expansion of NKT cells from HIV-1 infected patients. PBMCs from 3 groups (low viral load, medium viral load, and high viral load) of HIV-1 infected, antiviral therapy naïve patients were expanded in vitro by co-culture with alpha-galactosylceramide-loaded DCs. NKT cells, expanded from each ml of peripheral blood, were counted on days 0, 14, and 28 of co-culture. On day 14 of co-culture, NKT cells were re-stimulated by alpha-galactosylceramide/DCs. Symbols indicate various study subjects. Arrows indicate stimulation by alpha-galactosylceramide/DC. PBMCs from healthy individuals were used as control.

We observed no (subject HVL2) or very little expansion of NKT cells in patients with high viral load (i.e., those with Advanced AIDS) after 2 rounds of alpha-galactosylceramide-/DC stimulation.

### Alpha-galactosylceramide/DC triggered cytokine release by expanded NKT cells

We further tried to determine whether NKT cells expanded in HIV-1 infected patients were functionally capable of producing cytokines, upon being stimulated by alpha-galactosylceramide-loaded DCs. We used intracellular cytokine staining (ICS) technology to measure the percentage of NKT cells that produced cytokines. We generated a sufficient number of NKT cells from 3 low-viral-load, 3 medium-viral-load, and 4 healthy individuals for ICS experiments. We could not generate a sufficient number of NKT cells from any of the 5 high-viral-load (advanced AIDS) patients.

Figure 2 shows that, on average we observed 59% of expanded NKT cells produced IFN-γ in healthy individuals, 50% in medium-viral-load patients, and 70% in low-viral-load patients. We observed 65% of TNF-α producing NKT cells in healthy individuals, 58% in medium-viral-load patients, and 70% in low-viral-load patients. In addition to IFN-γ and TNF-α, 2 cytokines with direct antiviral effects, we also measured IL4, a Th2 cytokine with immunoregulatory functions. On average, we observed 5.8% of IL4 producing NKT cells in healthy individuals, 13.8% in medium-viral-load patients, and 4.1% in low-viral-load patients. Statistic analysis shows that the IL4 production of NKT cells expanded from medium-viral load patients is significantly higher than healthy control and low-viral-load groups (p<0.05).

**Figure 2.**
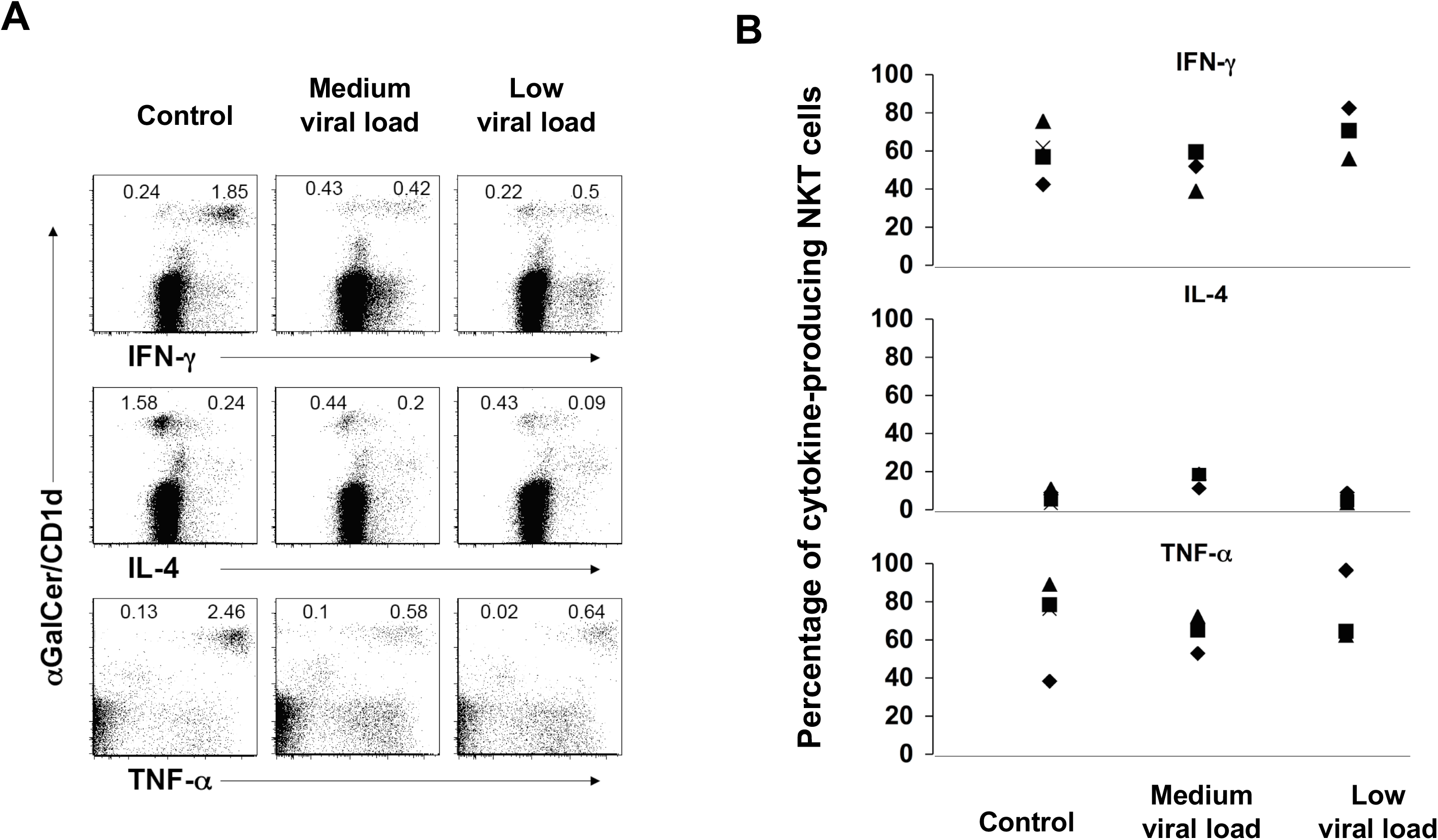
Cytokine production by expanded NKT cells from HIV-1 infected patients. Expanded NKT cells from HIV-1 infected patients were stimulated for 12 hours by alpha-galactosylceramide/DC in presence of Golgi-plug, and intracellular cytokines were stained by antibodies toward IFN-γ, IL4, and TNF-αrespectively. A. Profiles of NKT cells as stained by PBS57/CD1d tetramer and their expression of cytokines; B. Percentage of cytokine positive NKT cells in healthy control, medium viral load, and low viral load groups. Symbols indicate various study subjects. Only NKT cells from patients with medium and low viral loads could be studied. No sufficient number of NKT cells could be expanded from patients with high viral load (advanced AIDS patients).

## Discussion

Previous studies found that the number of NKT cells in HIV-1 infected patients is lower than that in healthy subjects (3–5). This reduction could not be corrected when peripheral blood NKT cells were incubated with alpha-galactosylceramide (9), the most potent ligand for NKT cells discovered so far. However, soluble alpha-galactosylceramide is not an efficient agent for NKT cell stimulation (15–18). Thus, we designed a study using monocyte derived DCs as antigen-presenting cells. Since CD1d, the antigen-presenting molecule for alpha-galactosylceramide, is nonpolymorphic in humans (1, 2), allogeneic DCs can efficiently present alpha-galactosylceramide, and provide other necessary costimulatory molecules and cytokines, such as CD80, CD86, CD40, IL12, and OX40L (21). A pitfall for this design is that allogenic DCs prepared from various donors, or in different batches, may express different levels of CD1d and costimulatory molecules. Therefore, we generated a large batch of DCs from buffy coat (up to 40 million DCs per donor per preparation) and froze them in 1 million-cell aliquots. We have used same batch of aliquots of DCs for our experiments on NKT cell expansion and cytokine production.

Our results indicate that when NKT cells in HIV-1 infected patients with low and medium viral loads are stimulated by potent ligands presented by DCs, they have the potential to expand. The magnitude of expansion in low-viral-load patients was similar to that in healthy individuals after 2 rounds of stimulation (28 days in culture), while medium-viral-load patients showed a decrease in expansion upon the second round of stimulation. These data suggest that in medium-viral-load patients, NKT cells may enter an early stage of being functionally exhausted, with decreased potential for responding to repeated alpha-galactosylceramide/DC stimulation. While in the 5 high-viral-load (advanced AIDS) patients, we found no NKT cell expansion, which may have been caused by both the lack of NKT cells and their functional exhaustion in these patients.

To further investigate the function of in vitro expanded NKT cells, we performed a cytokine production assay by stimulating NKT cells with alpha-galactosylceramide/DC. This assay measures the accumulation of intracellular cytokines within the first 12 hours upon alpha-galactosylceramide/DC stimulation. In low-viral-load patients, we observed similar or slightly higher IFN-γ (70%) and TNF-α (70%) production, as compared to that in healthy individuals (59% and 65% respectively). The levels of both cytokines were slightly lower in medium-viral-load patient group (50% and 58% respectively), although this result was not statistically significant. It is also note-worthy that a Th2 cytokine, IL4, recognized as an indicator for Th2-differentiation of NKT cells (1–2), was more abundantly produced in the medium-viral-load patients than healthy control group (p<0.05). Unfortunately, we could not generate a sufficient number of NKT cells from high-viral-load (advanced AIDS) patients to perform functional studies of their cytokine production. The mechanisms by which NKT cells lose their potential to expand and produce cytokines when viral load increases remain to be studied.

In conclusion, this study showed the previously underestimated potential of NKT cells as targets for functional reconstitution in HIV-1 infected patients with low and medium viral loads, before entering the advanced (AIDS) stage of disease.

## Authorship Contribution

LMB, HL, RCA, KJS, and DZ designed research; PT, PN, HH, BN, SJB, and DZ performed research; PT, PN, HL, JZ, RCA, KJS, and DZ analyzed and interpreted data; and DZ wrote the manuscript.

## Acknowledgement

We thank Luis Vence, Laszlo Radvanyi, Bindu Pappu, Markeda L Wade, and Kathryn B Carnes for support and discussion, Xiaoying Yu, Claudia Kozinetz, and Baylor-UTHouston CFAR Analysis and Design CORE for statistic analysis. DZ was supported by NIH grants R01 AI079232.

